# The variation among sites of protein structure divergence is shaped by mutation and scaled by selection

**DOI:** 10.1101/2020.07.10.196998

**Authors:** María Laura Marcos, Julian Echave

## Abstract

Protein structures do not evolve uniformly, but the degree of structure divergence varies among sites. The resulting site-dependent structure divergence patterns emerge from a process that involves mutation and selection, which may both, in principle, influence the emergent pattern. In contrast with sequence divergence patterns, which are known to be mainly determined by selection, the relative contributions of mutation and selection to structure divergence patterns is unclear. Here, studying 6 protein families with a mechanistic biophysical model of protein evolution, we untangle the effects of mutation and selection. We found that even in the absence of selection, structure divergence varies from site to site because the mutational sensitivity is not uniform. Selection scales the profile, increasing its amplitude, without changing its shape. This scaling effect follows from the similarity between mutational sensitivity and sequence variability profiles.

## 1. Introduction

Patterns of evolutionary variability inform on the processes that shape them. The evolutionary process has two main components: mutation, which originates new mutant genotypes, and selection, which determines the likelihood of mutants becoming lost or fixed [1]. At genotype level selection is usually the main force shaping emergent patterns. In contrast, at phenotype level even completely random unselected mutations may result in biased phenotypic variation governed not by selection, but by biases of the developmental process that maps genotypes onto phenotypes [2, 3, 4, 5, 6]. Therefore, when looking for the forces shaping observed patterns of evolutionary variation, it is important to remember that these patterns result, in general, from the combination of the effects of mutation and selection on phenotypes.

Here, we are interested in protein evolution, which results in the divergence of protein amino-acid sequences and their structures. A protein’s sequence and structure can be thought of as its genotype and phenotype, respectively, with protein folding, which maps the sequence onto the structure, playing the role of development. Most studies of patterns of protein divergence within protein families have focused on sequence divergence. In particular, much research has aimed at finding the biological and physical causes underlying the observed variation of sequence conservation among protein sites [7, 8, 9, 10, 11]. The main lesson is that the sequence divergence patterns are mostly governed by selection for stability [12, 13].

We know much less about patterns of structure divergence and their causes. Classic studies focused on overall structure divergence and its relation with sequence divergence and showed that structures evolve more slowly than sequences [14, 15, 16]. More recently, the focus has shifted to studying in more detail the directions of evolutionary deformation within protein families [17, 18, 19, 20, 21, 22]. In analogy with sequence patterns, some studies have tried selectionist explanations of structure patterns [17, 23]. However, other studies proposed that selection has no effect on structure patterns, which would be purely determined by mutational sensitivity [18, 19]. In such mutationist view, features of protein structure would be more variable or conserved according to whether they are more sensitive or insensitive with respect to mutations.

Despite the previous purely mutationist explanation of structure divergence patterns, it is still surprising that selection, all-important at sequence level, leaves no trace at structure level. Without selection, all protein sites would diverge with the same rate of amino acid substitutions per unit of time. Because of selection, different sites evolve at different rates. Selection being so evident in such variation among sites of of sequence divergence rates, here we hypothesised that in addition to mutational sensitivity, selection should also leave some trace in the variation among sites of the degree of structure divergence. If this was the case, site-dependent structure divergence profiles would be shaped by a combination of mutational sensitivity and natural selection.

To test the previous hypothesis, we dissected the relative effects of mutational sensitivity and selection on structure divergence. To this end, we derived a biophysical model of protein structure evolution, we validated the model by comparison with observed data, and we studied the differential effect of selection on structure divergence patterns by comparing patterns predicted under varying degrees of selection. In the following sections we describe methods in detail, present and discuss the results, draw the main conclusions and propose some directions of future research.

## 2. Methods

### 2.1. Structure divergence profiles: RMSD

To measure structure divergence patterns, we used `RMSD` profiles. Given a set of aligned and structurally superimposed homologous proteins, Root Mean Square profiles, `RMSD` = (`RMSD`_1_, `RMSD`_2_, …), were calculated as follows. First, we picked one member of the set, `ref`, as reference. Then, we obtained the pair alignments between `ref` and each other member of the set `p`. Next, we calculated the vectors 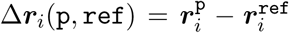, the difference between the position vectors of the alpha carbons of site *i* of `ref` and its homologous site of `p`. Finally, we obtained

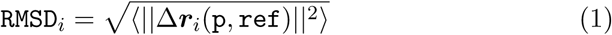

where ||.||^2^ is the squared Euclidean norm and *< … >* represents the average over `p` − `ref` pair alignments. Since we were not interested in absolute `RMSD` values but in their relative variation among sites, we normalized site scores:

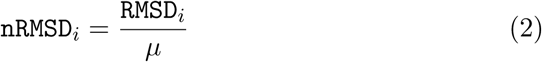

where *μ* is the average of `RMSD` over sites. Since we use `nRMSD` everywhere, for simplicity, we drop the *n* and use `RMSD` to denote normalized scores.

### 2.2. Data sets and RMSD^obs^

We studied 6 protein families chosen from the HOMSTRAD database of structurally aligned homologous protein families. HOMSTRAD is a database of carefully manually curated aligned and superimposed protein domains, often used as benchmark for automatic structure alignment algorithms [24, 25]. We picked families with at least 15 members, avoiding (1) families whose sequences were too similar,) which results in flat uninformative `RMSD` profiles, (2) families with proteins larger than 300 sites, which computational simulations too costly, and (3) families with proteins smaller than 50 sites, too small to be modelled accurately using the sort of Elastic Network Models used here. The families chosen, and some key properties are summarized in Table 1.

**Table 1:** Protein families

| family | $N_{\text{prot}}$ | $\langle \text{Id}\% \rangle$ | ref pdb | $N_{\text{sites}}$ | SCOP |
| --- | --- | --- | --- | --- | --- |
| Serine proteases | 27 | 38 | 1mct | 223 | $\beta$ |
| Fatty acid binding proteins | 17 | 36 | 1hmt | 131 | $\beta$ |
| Globins | 41 | 32 | 1a4f,B | 146 | $\alpha$ |
| Phospholipases A2 | 18 | 49 | 1jia,A | 122 | $\alpha$ |
| RNA recognition motif | 20 | 23 | 1fxl | 82 | $\alpha/\beta$ |
| Src homology 3 domain | 20 | 32 | 1lck,A | 54 | small |

Columns show, in order, family name, brief family id, number of members, average % sequence identity, reference protein (its pdb id and chain), number of sites of the reference, and structure class according to the SCOP classification [26].

`RMSD`^obs^, the *observed profiles*, are the `RMSD` profiles calculated using the HOMSTRAD alignments. For each family, we chose as reference protein `ref` the protein whose structure is closer to the average family structure (Table 1). Then, we calculated the structure divergence profile `RMSD`^obs^ using this reference, as explained in section 2.1.

### 2.3. Mutation-Selection Model and RMSD^MSM^

The Mutation Selection Model, `MSM`, is a biophysical model of protein evolution that combines previous models used to study structure divergence [18, 19] and sequence divergence [27]. Briefly, proteins are represented by Elastic Network Models, mutations as perturbations of the network contacts, selection using a stability-based fixation probability, and simulations are performed by repeating mutation-selection steps. In this section, we describe in some detail each model component.

#### 2.3.1. Elastic Network Model

We used the simplest Elastic Network Model (ENM), the so-called Anisotropic Network Model, ANM [28]. The ANM represents sites as nodes placed at the sites’ *C*_*α*_ positions and joins nodes using springs of identical spring constant *k*, if they are within a cut-off distance *R*_0_ of each other. For the wild-type protein at a given point of time, ANM represents the protein’s potential energy as:

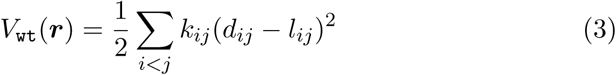

where *d*_*ij*_ is the distance between nodes *i* and *j* in a given protein conformation ***r***, and *l*_*ij*_ is the length of spring *i* − *j*. All models developed so far use 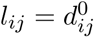, where 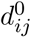 is the distance between *i* and *j* in the native conformation 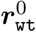. This guarantees that this conformation is at the minimum of the energy well. The force constants are given by:

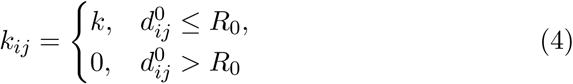

Therefore, ANM has two parameters, *R*_0_ and *k*. In general, results are quite insensitive to the precise values in the range 10 ≤ *R*_0_ ≤ 15. Here we used *R*_0_ = 12.5*Å*. The parameter *k* has no effect on normalized `RMSD` values, of interest here, thus we arbitrarily set *k* = 1.

For proteins, the most common use of ENMs is to analyse protein motions. For instance, normal modes can be obtained by diagonalization of the matrix of second derivatives of *V*_wt_(***r***), the Hessian matrix **K**. This matrix can also be used to calculate the variance-covariance matrix **C**:

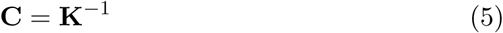

(Rigorously, **K** is singular and has no inverse, thus the pseudo-inverse is used instead, see e.g. [19].) **C** can be partitioned into blocks **C**_*ij*_, where *i* and *j* designate sites. Off-diagonal blocks describe correlated motions and diagonal blocks characterize individual fluctuations. For example, the flexibility of site *i*, measured by its Root Mean Square Fluctuation, `RMSF`, is given by:

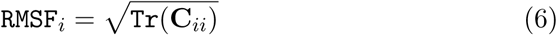

where `Tr` is the matrix trace operation.

#### 2.3.2. Mutation

We represent point mutations as small perturbations of the springs connecting the mutated site to its neighbours [18, 19]. Specifically, a mutation is modelled as:

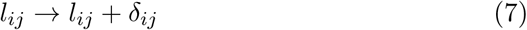

where *δ*_*ij*_ are the spring-length perturbations which are zero except for contacts of the mutated site, for which they are independently chosen from identical normal distributions:

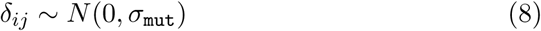

We arbitrarily set *σ*_mut_ = 1 in the present study, because this parameter has no effect on the normalized `RMSD` profiles, which are our focus here.

The mutant’s energy function is given by:

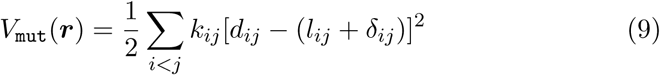

The mutant’s native conformation, 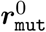 is the ***r*** that minimizes *V*_mut_(***r***). It may be calculated using:

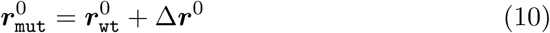

where Δ***r***^0^ is the deformation that results from the mutation and can be calculated using:

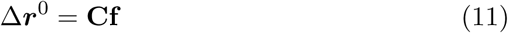

where **C** is the variance-covariance matrix given by Eq. 5 and **f** is a “force” vector that can be calculated from the perturbations *δ*_*ij*_ of Eq. 7 [18, 19].

#### 2.3.3. Selection

`MSM` is an origination-fixation model [1]: at any given time the population consists of a single genotype, the current wild type, and evolution results from repeated mutation-fixation steps consisting of the origination of a mutant genotype by mutation, which either disappears or gets fixed, replacing the wild-type, according the value of a fixation probability function, which represents natural selection.

Here, natural selection is modelled by the following fixation probability function [27]:

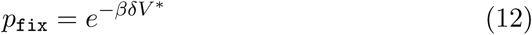

where *β* is a parameter that allows tuning selection pressure against destabilization and *δV* ^∗^ is the energy difference between the mutant and the wild type when both are at the active conformation, which is assumed to be the native wild-type equilibrium structure:

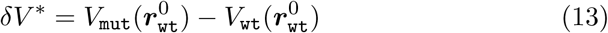

This model of selection was derived in detail in [27]. Noteworthy, an exponential fixation probability such as Eq. 12 is related to a fitness function that is a “step function” [8, 29]. The interested reader could find more details in the cited references.

#### 2.3.4. Simulation

To generate an evolutionary path of protein structures ***r***_0_, ***r***_1_, …, ***r***_*n*_, we need to specify the initial structure and the model selection parameter *β*. The path starts at ***r***_*t*_ = ***r***_0_. At time-step *t*, a mutation is introduced in ***r***_*t*_ by picking a protein site randomly and perturbing all its contacts to obtain a trial mutant, the energy *δV* ^∗^ and fixation probability *p*_fix_ are calculated, and the mutation is accepted if a number between 0 and 1 picked from a uniform distribution is smaller than *p*_fix_. If the mutation is fixed, ***r***_mut_ is calculated and ***r***_*t*+1_ = ***r***_mut_ is set. This mutation-selection step is repeated until *t* = *n* is reached.

The previous process can be used to generate any desired phylogenetic tree. In this work, we simulated star trees, which are sets of independent paths of the same length divergent from the same ancestral structure ***r***_0_. To generate a star tree, therefore, model input is selection parameter *β*, common ancestor ***r***_0_, lineage length *n* = *n*_subs_, and number of lineages *n*`paths` = *n*`lineages`.

#### 2.3.5. RMSD^MSM^

`RMSD`^MSM^ were calculated from aligned proteins obtained from `MSM` simulations. For each family studied, we chose the member closest in structure to the family average as reference protein (Table 1). Then we generated simulated datasets using `MSM` for increasing selection pressures *β*. We used *n*_branches_ = 100 and *n*_subs_ = (100 − `Id`%)/100, where *Id*% is the average sequence identity percent of the HOMSTRAD family. Using these simulated datasets, we calculated the site-dependent structure divergence profiles `RMSD`^MSM^, as explained in section 2.1.

#### 2.3.6. Acceptance rate: ω

In addition to structure divergence degrees, measured by `RMSD`, we consider also sequence divergence degrees, which are conveniently measured using *ω*, the so-called acceptance rate, which is the average number of fixations per mutation. At protein level, *ω* can be calculated using:

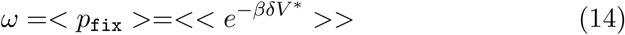

where the double-averaging symbol *<< … >>* is used to denote a double averaging, over sites and mutations. At site level, *ω*_*i*_ is given by

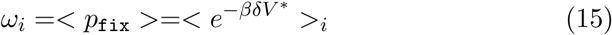

with *< … >*_*i*_ denoting an average over mutations at site *i*:

*ω* is a convenient measure of selection pressure. If *β* = 0 there is no selection: all mutations are accepted, *p*_fix_ = 1, and *ω* = 1. On the other hand, as the selection pressure parameter *β* increases, *ω* decreases.

### 2.4. Mutation Model and RMSD^MM^

We use the Mutation Model, `MM`, as a control to quantify the effect of adding selection over the purely mutational effect. `MM` is the special case of `MSM` with *β* = 0 (*ω* = 1) in which all mutations are fixed and thus selection does not operate. Therefore, `RMSD`^MM^ = `RMSD`^MSM^(*β* = 0), calculated as described in the previous section. Since in this case the only source of variation among sites is the variable sensitivity of the structure’s response to mutations, we call `RMSD`^MM^ the *mutational sensitivity*.

## 3. Results and discussion

We studied the effects of mutation and selection on site-dependent structure divergence patterns for the 6 protein families of Table 1. To this end, we compared three types of profile: `RMSD`^MSM^, `RMSD`^MM^, and `RMSD`^obs^. An `RMSD` profile is a vector `RMSD` = (`RMSD`_1_, `RMSD`_2_, …), where `RMSD`_*i*_ quantifies the relative structure divergence of site *i*. `RMSD`^MSM^ are profiles predicted by the Mutation Selection Model, `MSM`, which simulates evolutionary divergence under the effect of mutational deformations and selection on protein stability. `RMSD`^MM^ are profiles predicted by the Mutation Model, `MM`, which is the special case of `MSM` without selection (and, therefore, represents the structure response of the protein structure to unselected random mutations, which is why we refer to `RMSD`^MM^ as the *mutational sensitivity*). `RMSD`^obs^ are the observed profiles, calculated from the multiple alignments of actual family members.

For clarity, we used one example case, serine proteases, for the figures shown in the main document. Results for other families are similar, as can be seen in supplementary figures (section 4.)

### 3.1. Model predictions

We start by analysing the effect of varying selection pressure on model predictions. We run `MSM` simulations for all families and various values of the selection parameter *β*. Then, for each family and each *β* value, we calculate the `RMSD`^MSM^ profile and the average fixation probability *ω*, which is a convenient measure of selection pressure.

`RMSD`^MSM^ profiles for serine proteases are shown in Figure 1. Clearly, structure divergence varies among sites even in the absence of selection (*ω* = 1). As selection pressure increases (*ω* decreases), `RMSD`^MSM^ profiles become more pronounced. However, while increasing selection makes peaks higher and wells deeper, selection does not affect the positions of peaks and wells along the sequence, which is why all `RMSD`^MSM^ profiles look similar to the mutational sensitivity profile `RMSD`^MM^.

**Figure 1:**
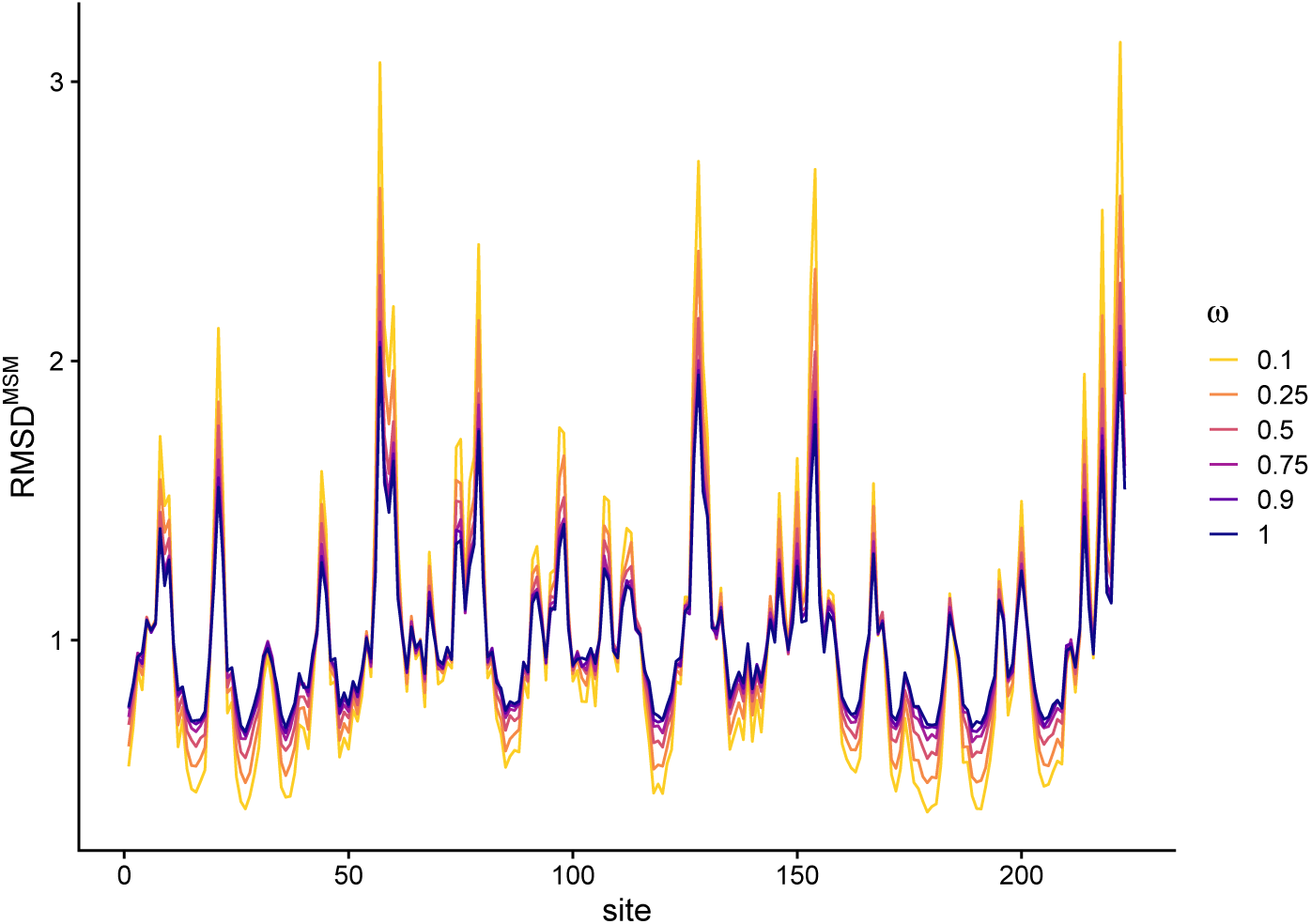
Predicted site-dependent structure divergence profiles for serine proteases. `RMSD`^MSM^ is the site-dependent profile of Root Mean Square Deviations calculated from sets of proteins simulated with the Mutation Selection model `MSM`. `RMSD` values are normalized so that their average over sites is 1. `site` is the site of the protein used as initial condition of simulations. *ω* is the average probability of accepting mutations. As (negative) selection pressure increases, *ω* decreases. The maximum is *ω* = 1, which is the no-selection case in which all mutations are accepted. We call this special case the Mutational Model, and we denote the corresponding profile `RMSD`^MM^ (`RMSD`^MM^ = `RMSD`^MSM^(*ω* = 1)). It is important to note that `RMSD`^MM^ is not uniform: even in the absence of selection structure divergence varies among sites, because the sensitivity of the structure to random mutations varies among sites. Therefore, `RMSD`^MM^ represents the *mutational sensitivity*. Selection makes the profiles more pronounced, but does not seem to affect their shape.

More quantitatively, we find that `RMSD`^MSM^ increases linearly with `RMSD`^MM^ (Figure 2**a**), obeying:

**Figure 2:**
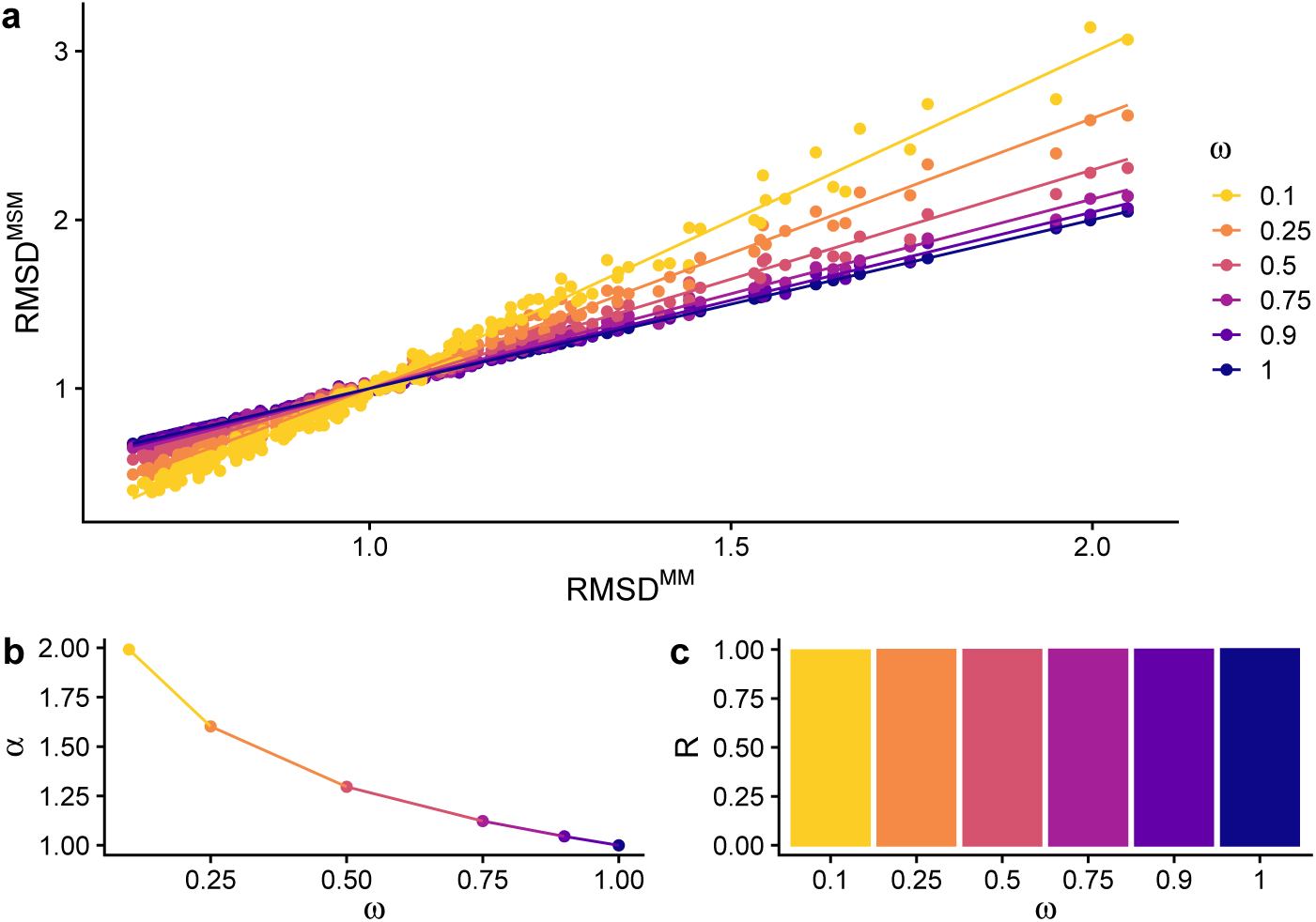
structure divergence is proportional to mutational sensitivity. The results shown are for serine proteases. **a**: structure divergence, `RMSD`^MSM^ depends linearly on mutational sensitivity, `RMSD`^MM^. Points represent sites and lines were obtained fitting the points using Eq.16: `RMSD`^MSM^ = 1 + *α*(`RMSD`^MM^ − 1). **b:** Increasing selection pressure (decreasing *ω*) increases the slope of the `RMSD`^MSM^ vs. `RMSD`^MM^ lines, *α*. Therefore, selection linearly scales the profile making it more pronounced. **c:** Pearson correlation coefficient, *R*, between `RMSD`^MSM^ and `RMSD`^MM^. That *R ≈* 1 for all *ω* values, means that while selection scales the `RMSD`^MSM^ profile, it does not affect its shape, which is determined by the mutational sensitivity `RMSD`^MM^.

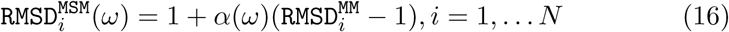

where *i* is the site index and *N* the number of sites, *ω* is the average acceptance probability, and *α* is the slope of the linear dependence.

According to Eq. 16, any variation among sites of `RMSD`^MSM^ is due to the mutational sensitivity `RMSD`^MM^, which does not depend on selection. Selection affects only the site-independent slope parameter *α*, which has a minimum *α* = 1 for *ω* = 1 and increases with decreasing *ω* (Figure 2**b**). Therefore, Eq. 16 accounts for the observed similarity between `RMSD`^MSM^ and `RMSD`^MM^ profiles and for the amplifying effect of selection.

Note that according to Eq. 16, `RMSD`^MSM^ and `RMSD`^MM^ are identical, modulo a linear transformation, so that, in this very sense, they can be said to have the same shape. On the other hand, the increase of the parameter *α* with selection pressure (Figure 2**b** and Figure 2**c**), accounts for the scaling of `RMSD`^MSM^ profiles (Figure 2**a**). Therefore, we can say that the variation among sites of structure divergence is shaped by mutational sensitivity and scaled by selection.

### 3.2. Predictions vs observations

To test the previous predictions, we assessed whether Eq. 16 fits observed profiles, `RMSD`^obs^. The model-vs-data fit for serine proteases is visualized in Figure 3. In this case, the relationship between the profiles compared is `RMSD`^obs^ ≈ `RMSD`^MSM^ = 1+1.9(`RMSD`^MM^ − 1) (Table 2). Thus, the observed profile `RMSD`^obs^ is similar to the `MSM` profile `RMSD`^MSM^ and both are similar to, but more pronounced than the mutational sensitivity profile `RMSD`^MM^ (Figure 3**a** and Figure 3**b**). Accordingly, `RMSD`^obs^ increases with `RMSD`^MM^ more steeply than expected in the absence of selection (Figure 3**c**).

**Table 2:** Best linear fit to observed profiles, `RMSD`^MSM^ = 1 + *α*(`RMSD`^MM^ − 1).

| family | $R$ | $\sigma$ | $\alpha \pm \text{se}$ | $\omega \pm \text{se}$ |
| --- | --- | --- | --- | --- |
| Serin proteases | 0.70 | 0.50 | $1.90 \pm 0.13$ | $0.13 \pm 0.05$ |
| Fatty acid binding proteins | 0.74 | 0.32 | $1.98 \pm 0.16$ | $0.04 \pm 0.1$ |
| Globins | 0.62 | 0.36 | $1.50 \pm 0.16$ | $0.28 \pm 0.13$ |
| Phospholipases A2 | 0.72 | 0.40 | $1.37 \pm 0.12$ | $0.34 \pm 0.14$ |
| RNA recognition motif | 0.80 | 0.43 | $1.33 \pm 0.11$ | $0.32 \pm 0.13$ |
| Src homology 3 domain | 0.77 | 0.39 | $1.32 \pm 0.15$ | $0.26 \pm 0.16$ |

$R$ is the Pearson correlation coefficient between $\text{RMSD}^{\text{obs}}$ and $\text{RMSD}^{\text{MSM}}$ ; $\sigma$ is the standard deviation of the residuals; $\alpha$ is the slope estimate of the best model-data fit and $\text{se}$ its standard error; $\omega$ is the mutation acceptance rate predicted by the best MSM model and $\text{se}$ its standard error. Note that for all cases $\omega < 1$ , which means that selection does affect the observed structure divergence profiles.

**Figure 3:**
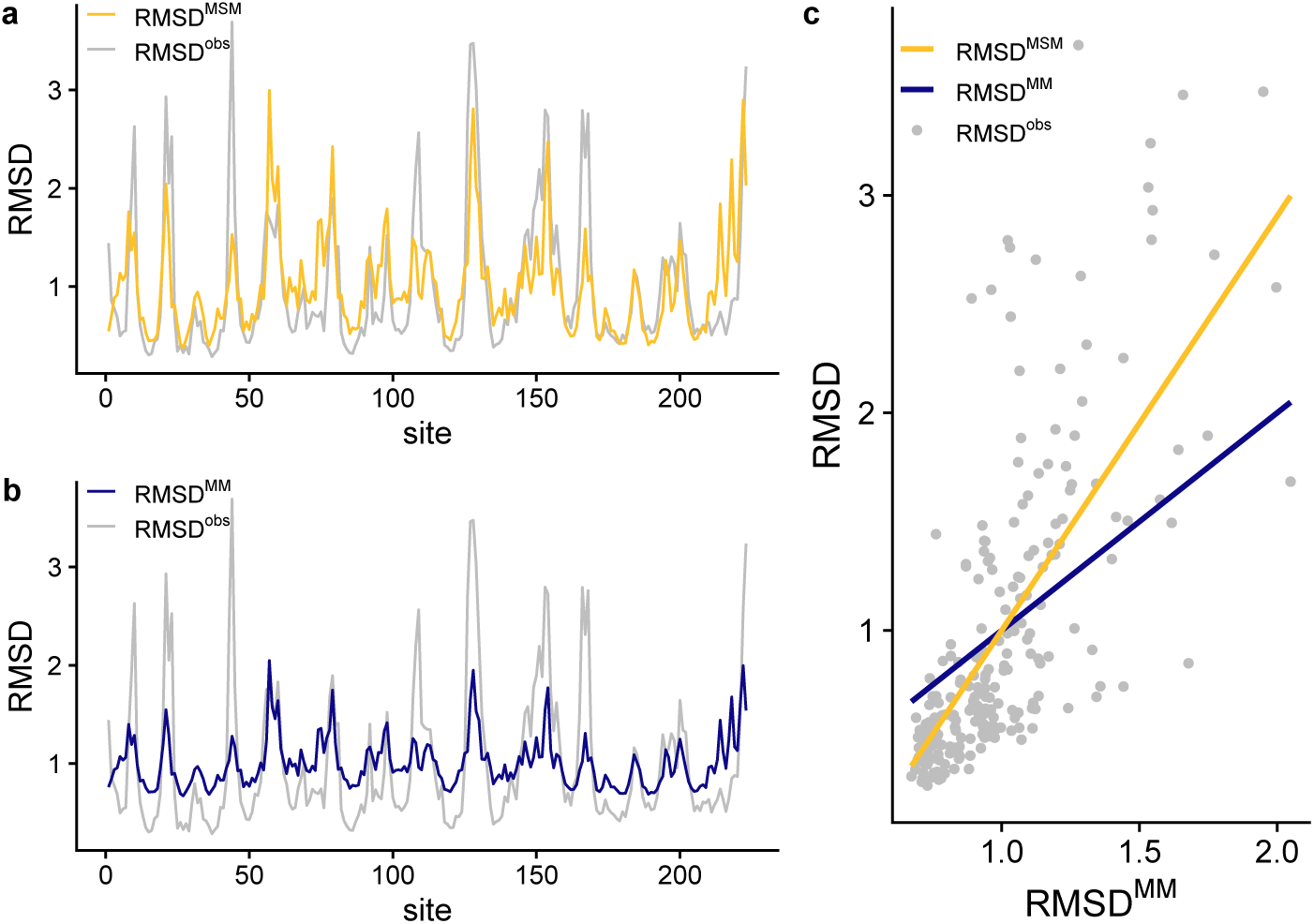
Predicted vs. observed structure divergence profiles for serine proteases. Three structure divergence profiles are shown: the observed profile `RMSD`^obs^, the predicted profile `RMSD`^MSM^ obtained by fitting Eq. 16 to observed data, and, as control, the mutational sensitivity profile RMSD^MM^. The differences between `RMSD`^MSM^ and `RMSD`^MM^ are due to selection. **a:** `RMSD`^obs^ vs. `RMSD`^MSM^; note that `RMSD`^obs^ is very similar to `RMSD`^MSM^. **b:** `RMSD`^obs^ vs. `RMSD`^MM^; both profiles are similar in shape but `RMSD`^obs^ has larger amplitude because of the amplifying effect of selection. **c:** structure divergence vs. mutational sensitivity. Note that the `RMSD`^MSM^ vs. `RMSD`^MM^ line does not superimpose with the *y* = *x* line, also shown. This is due to selection, which increases the slope, amplifying the variation of structure divergence among sites.

In general, Eq. 16, provides a very good fit to `RMSD`^obs^ for all families of this study, which validates the `MSM` model (Table 2). The Pearson correlation coefficient between `RMSD`^obs^ and `RMSD`^MM^ varies in the range 0.62 *≤ R ≤* 0.8, which shows that the mutational sensitivity is a major determinant of structure divergence. In addition, in all cases *α >* 1, consistent with the presence of negative selection, as modelled by the `MSM` model.

In summary, as predicted by the `MSM` model, observed profiles `RMSD`^obs^ are similar to the mutational sensitivity profile `RMSD`^MM^, but more pronounced, because they’re scaled by selection with a scaling parameter *α >* 1.

### 3.3. Dependence of RMSD on flexibility and packing

According to Eq. 11, mutational deformations depend on the protein’s variance-covariance matrix **C**. This matrix characterizes protein fluctuations (Eq. 6)), which suggests a possible relationship between structure divergence and flexibility. In addition, since flexibility is related to local packing [30], we would also expect a relationship between divergence and packing. Indeed, as we show in Figure 4, both `RMSD`^MSM^ and `RMSD`^MM^ increase with `RMSF` and decrease with `CN`. These dependencies should be seen as representations of a single gradient, because 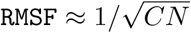 [30]. The gradient is in place even without selection, due to variation among sites of mutational sensitivity (`RMSD`^MM^). With selection (`RMSD`^MSM^), the gradient becomes steeper.

**Figure 4:**
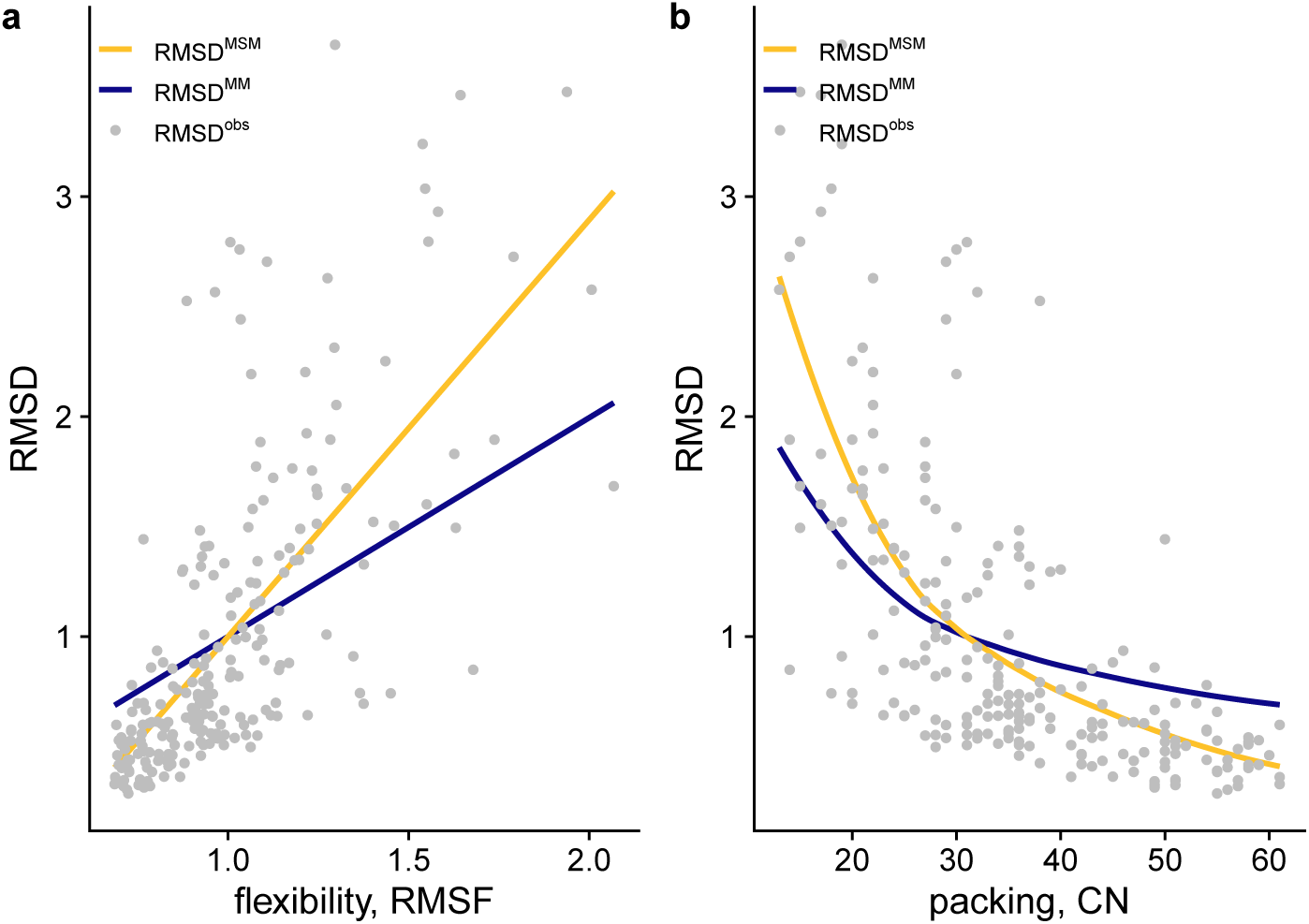
Dependence of RMSD on flexibility and packing for serine proteases. Comparison of observed site-specific structure divergence, `RMSD`^obs^, with predictions `RMSD`^MSM^ and `RMSD`^MM^. **a:** Increase of structure divergence with increasing site flexibility, measured by `RMSF` (the Root Mean Square Fluctuations predicted by the Anisotropic Network Model). **b:** Decrease of structure divergence with increasing local packing density, measured by the Contact Number CN (number of sites within a distance of 12.5 Å). The two dependencies are related because 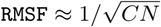, thus these two figures are alternative representations of a single gradient: an increase of structure divergence from the tightly packed and rigid protein core towards the loosely packed and flexible regions. Importantly, this gradient is evident even without selection; selection has a seemingly secondary effect of making it steeper without changing its shape.

### 3.4. Structure patterns vs. sequence patterns

So far, we have shown that `RMSD`^MSM^ increases linearly with mutational sensitivity `RMSD`^MM^ with a constant slope that depends on selection pressure (Eq. 16). Further, we showed that `RMSD`^MM^ increases with flexibility and decreases with packing, with selection making these gradients steeper. What is the mechanism that underpins such seemingly secondary role of selection?

To answer this question, we compare structure profiles with sequence profiles (Figure 5). Without selection, all sites evolve at the same rate: the sequence *ω* profile is flat. In contrast, even without selection, `RMSD`^MM^ varies among sites, as we know from previous sections. Because of selection, different sites evolve at different rates, and a site-dependent sequence profile emerges. At structure level, selection does not affect the shape of the `RMSD` profile, but just scales it. From the figure, it is obvious that mutational sensitivity and sequence profiles are very similar. Because of this similarity, selection does not change the shape of the `RMSD` profile but only its amplitude, which explains the scaling effect of selection described by Eq. 16.

**Figure 5:**
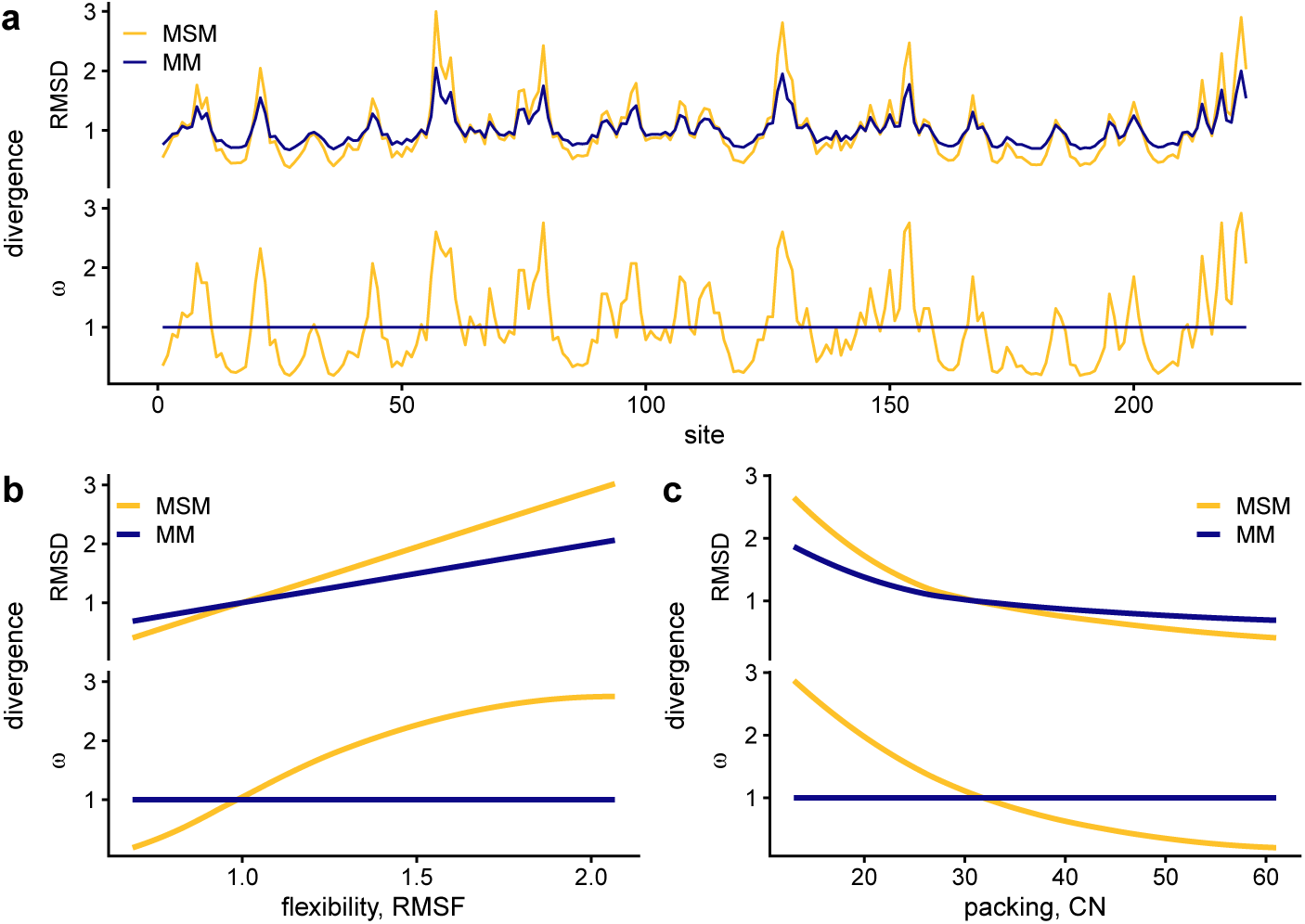
Comparison of structure and sequence divergence profiles. Structure divergence is measured by `RMSD` and sequence divergence by acceptance rates *ω*. Both are normalized so that their average over sites is 1. Predictions of the `MSM` model that best fits `RMSD`^obs^ observed profiles are compared with predictions of the purely mutational `MM` model. `RMSF` measures site flexibility, and the contact number `CN` measures local packing. **a**: Comparison of structure divergence profiles (top) and sequence divergence profiles (bottom). **b:** Comparison of `RMSD-vs-RMSF` and *ω*-`vs-RMSF` gradients. **c:** Comparison of `RMSD-vs-CN` and *ω*-`vs-CN` gradients. Note that in the absence of selection (`MM` model), sequence divergence is identical for all sites (flat profiles), but structure divergence varies among sites. With selection, the sequence divergence profile emerges and structure divergence becomes amplified. This amplification is due to `RMSD`^MSM^ resulting from a combination of two very similar profiles: the selection-independent mutational sensitivity profile, `RMSD`^MM^, and the selection-dependent sequence profile, *ω*^MSM^.

### 3.5. Comparison of different ENM models

All the previous results are based on the Anisotropic Network Model (ANM). This model, assigning the same spring force constant *k*_*ij*_ = *k* to all contacts, is the simplest possible elastic network model. From the very good agreement between predictions and observations (Table 2), it follows that in spite of its simplicity, the ANM-based `MSM` model captures the physics underlying the variation of structure divergence among sites. However, other presumably more realistic network models exist and it is worthwhile to test the effect of ENM model choice on `RMSD` predictions. To this end, we compared ANM results with three alternative elastic network models: MW, HCA, and pfANM. MW uses two different force constants, a larger one for contacts between covalently bound residues and a smaller one for noncovalent contacts[31]; HCA, the “harmonic alpha carbon” model, sets a large force constant for pairs with *d*_*ij*_ < 4.5*Å*, which includes the covalently bound pairs, and 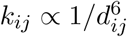 for other pairs [32]; pfANM, the “parameter-free Anisotropic Network Model”, uses 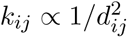 for all pairs [33].

From Figure 6, we see that the `RMSD`^MSM^ profiles predicted by different ENM models are similar to each other and similar to the observed profiles `RMSD`^obs^. More quantitatively, the goodness of fit between `RMSD`^MSM^ and `RMSD`^obs^ is similar for different models (Table 3). The reason for such model-independence is that evolutionary deformations are governed by the variance-covariance matrix **C** (see Eq. 11), which depends more on the elastic network topology, captured by all models, than on details of the interactions, which is why elastic network models work [34].

**Table 3:** Comparison of `RMSF` profiles predicted by different models.

| family | ANM | MW | HCA | pfANM |
| --- | --- | --- | --- | --- |
| Serin proteases | 0.70 | 0.63 | 0.67 | 0.69 |
| Fatty acid binding proteins | 0.73 | 0.48 | 0.70 | 0.66 |
| Globins | 0.62 | 0.51 | 0.59 | 0.58 |
| Phospholipases A2 | 0.73 | 0.65 | 0.72 | 0.72 |
| RNA recognition motif | 0.80 | 0.77 | 0.79 | 0.78 |
| Src homology 3 domain | 0.77 | 0.72 | 0.77 | 0.77 |

The table shows $R$ , the Pearson correlation coefficient between $\text{RMSD}^{\text{obs}}$ and $\text{RMSD}^{\text{MSM}}$ predicted by four different elastic network models: the anisotropic network model (ANM), Ming’s and Wall’s model (MW), the harmonic $C_\alpha$ model (HCA), and the parameter-free anisotropic network model (pfANM). Models are described in section 3.5.

**Figure 6:**
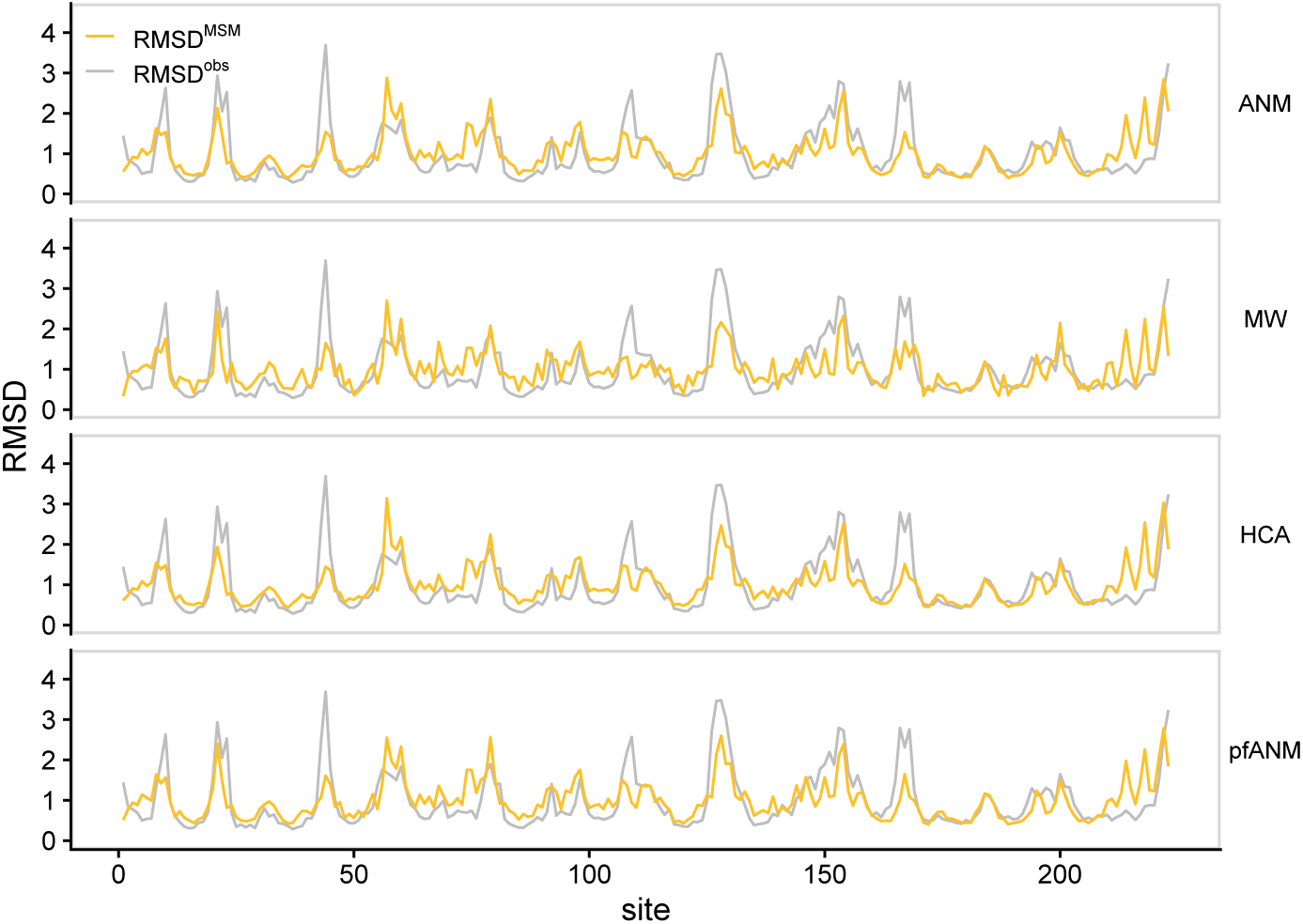
Dependence of predictions on ENM model choice, case of serine proteases. Comparison of `RMSD`^MSM^ predictions using four different elastic network models denoted ANM, MW, HCA, and pfANM (defined in section 3.5). The observed profile `RMSD`^obs^ is also shown. All models predict similar profiles.

## 4. Conclusion

To summarize, we studied the relative effects of mutational sensitivity and natural selection on the variation among sites of structure divergence in protein families. We found that both mutation and selection contribute to structure divergence. Specifically, the shape of the site-dependent structure divergence profile is determined by mutational sensitivity, and selection has a seemingly secondary effect of scaling the profile (Eq. 16). We also showed that mutational sensitivity increases with site flexibility and decreases with local packing and that, again, these gradients become steeper because of the scaling effect of selection. Finally, we proposed an explanation of this scaling effect of selection: the mutational sensitivity profile and the sequence variability profiles are similar, so that their combination does not affect the profile shape, but only its scale.

To finish, we mention some future research directions we find interesting. First, the most straightforward follow-up of this work would be to to perform a detailed comparative study of site-dependent patterns at sequence and structure levels, to look for subtle traces of selection in divergence patterns, beyond the general trend demonstrated here. Second, this study focused on stability-based selection, which is the main force shaping sequence divergence patterns, but selection on activity may also have strong effects, especially at and close to active sites. It would be interesting to see how this affects structure conservation profiles. Third, depending on the findings of the previous two lines, the difference between `RMSD`^MSM^ predictions and `RMSD`^obs^ observations could help detect functional sites. Finally, at present the `MSM` model does not consider insertions of deletions, which may limit its applicability to close homologues at the family level. Therefore, it would be worthwhile to extend the model to consider the effects of insertions and deletions in order to study more diverged structures.

## Acknowledgements

This work was supported by Consejo Nacional de Investigaciones Científicas y Técnicas (grant number PIP 112 201501 00385 CO) and by Agencia Nacional de Promoción Científica y Tecnológica (grant number PICT-2016-4209).

## Supplementary information

Supplementary figures analogous to the ones shown, but for other families, can be in `suppl.pdf`.

## Notes

### Competing Interest Statement

The authors have declared no competing interest.

