## Supplemental figures for "The variation among sites of protein structure divergence is shaped by mutation and scaled by selection"

For a detailed description of each figure, see caption of equivalent figure in the main document.

### Fatty acid binding proteins

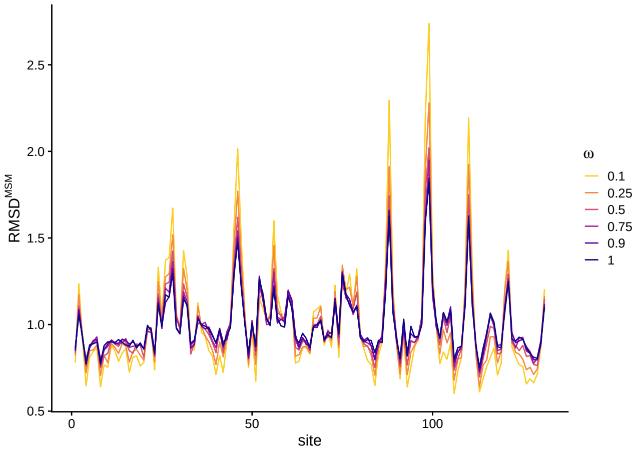


Figure S1: **Predicted site-dependent structure divergence profiles.**

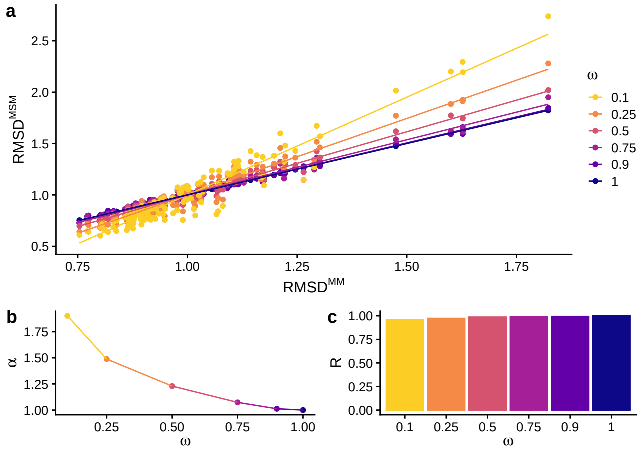


Figure S2: **Structure divergence is proportional to mutational sensitivity.**

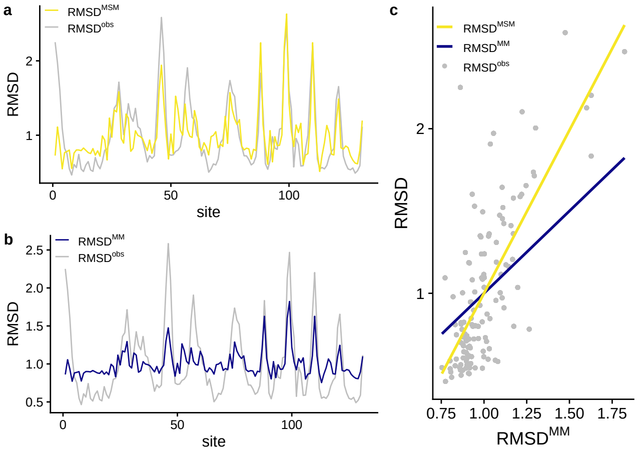


Figure S3: **Predicted vs. observed structure divergence profiles.**

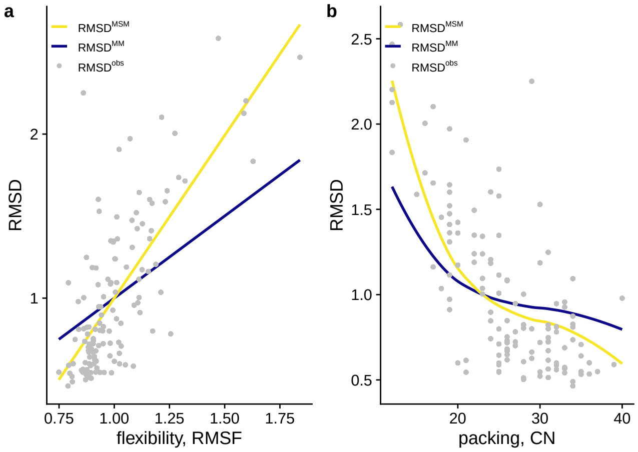


Figure S4: **Dependence of `RMSD` on flexibility and packing.**

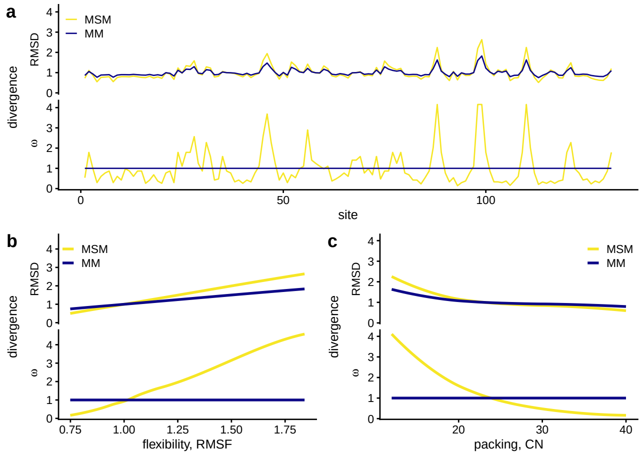


Figure S5: **Comparison of structure and sequence divergence profiles.**

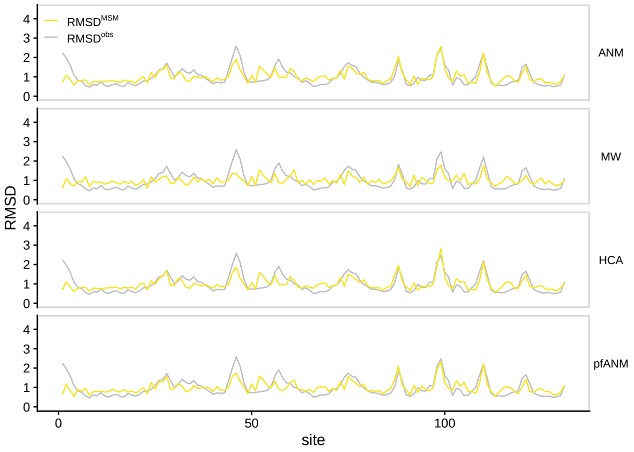


Figure S6: **Dependence of predictions on ENM model choice.**

### Globins

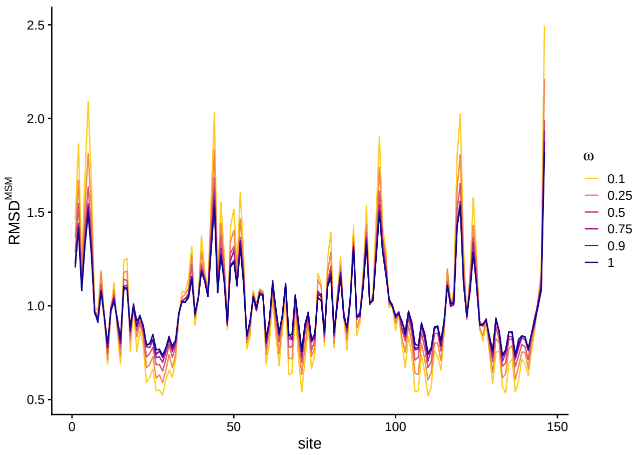


Figure S7: **Predicted site-dependent structure divergence profiles.**

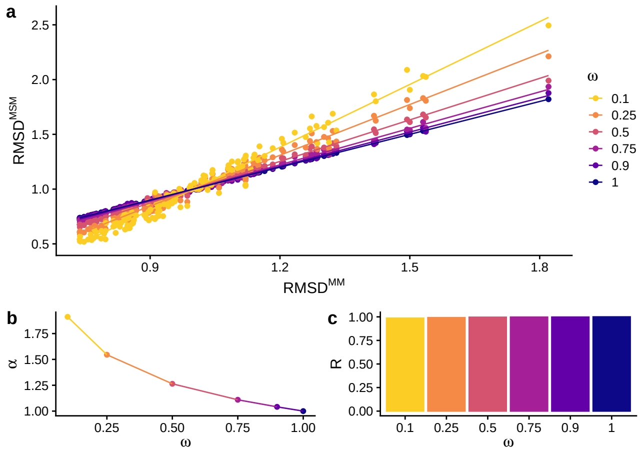


Figure S8: **Structure divergence is proportional to mutational sensitivity.**

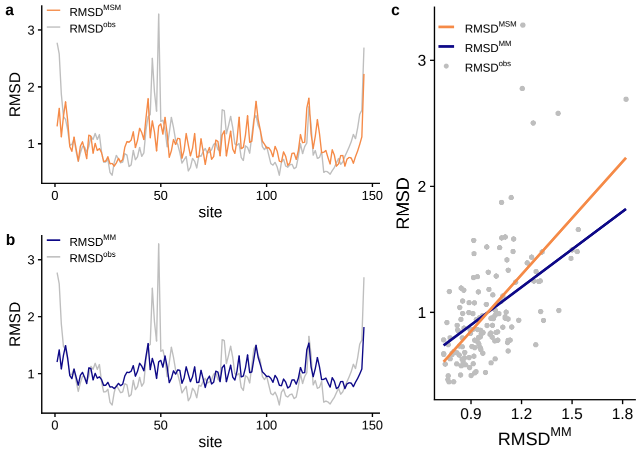


Figure S9: **Predicted vs. observed structure divergence profiles.**

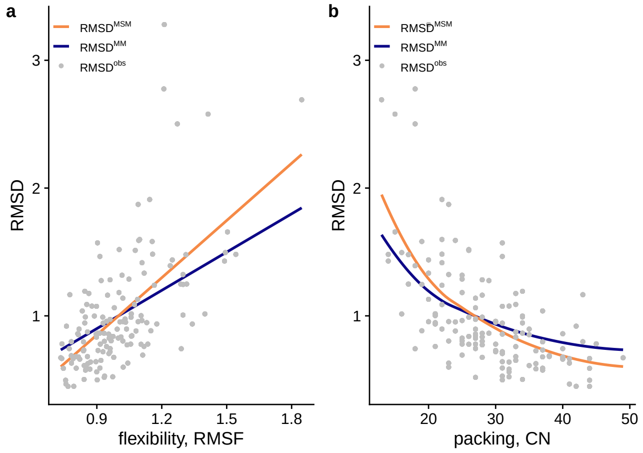


Figure S10: **Dependence of `RMSD` on flexibility and packing.**

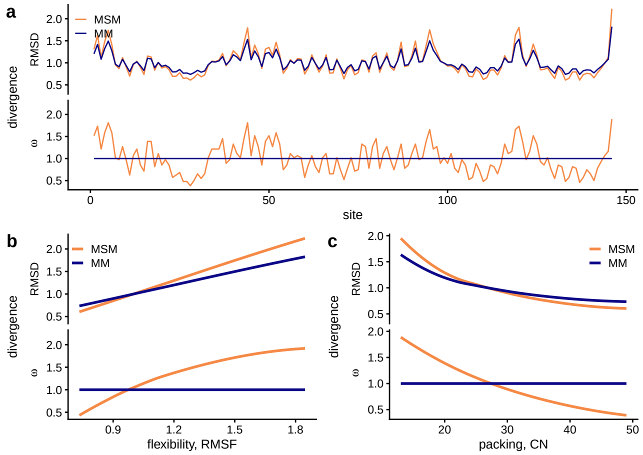


Figure S11: **Comparison of structure and sequence divergence profiles.**

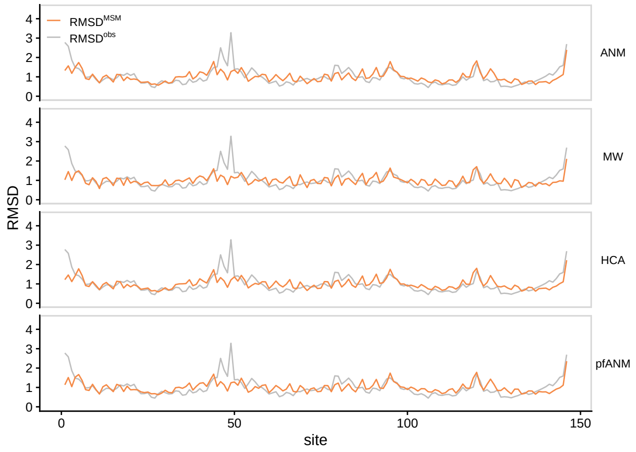


Figure S12: **Dependence of predictions on ENM model choice.**

### Phospholipases

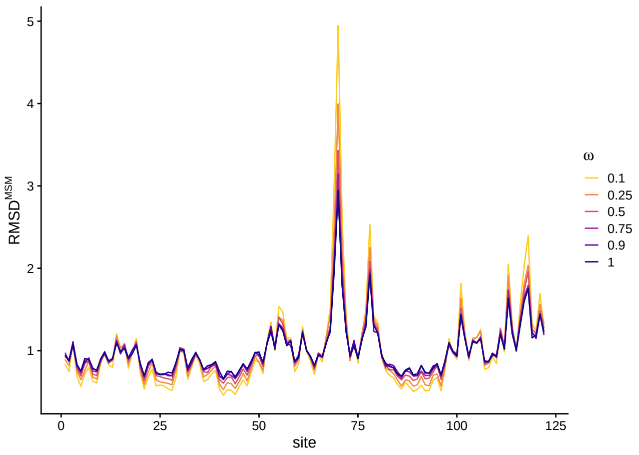


Figure S13: **Predicted site-dependent structure divergence profiles.**

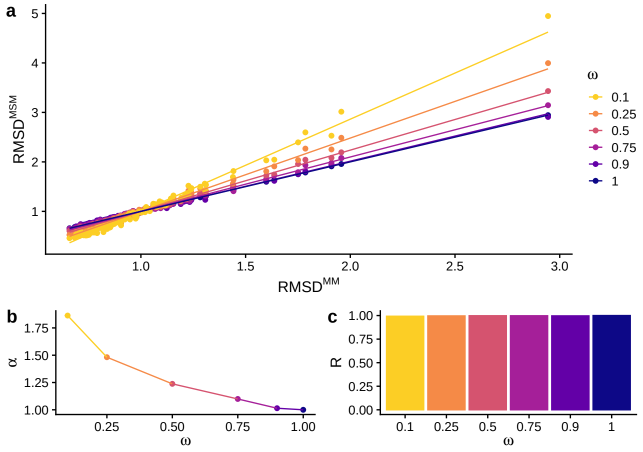


Figure S14: **Structure divergence is proportional to mutational sensitivity.**

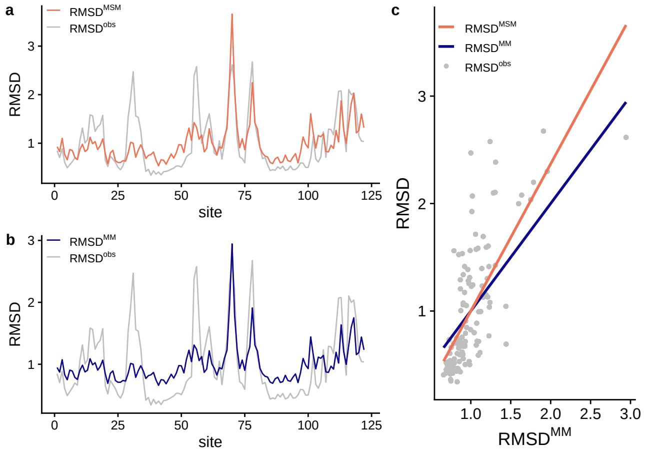


Figure S15: **Predicted vs. observed structure divergence profiles.**

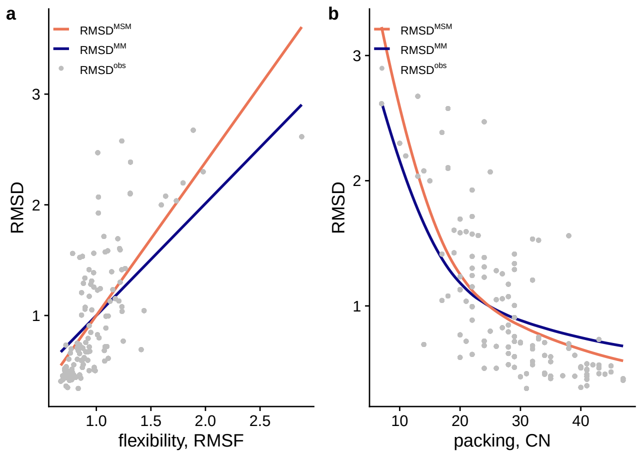


Figure S16: **Dependence of `RMSD` on flexibility and packing.**

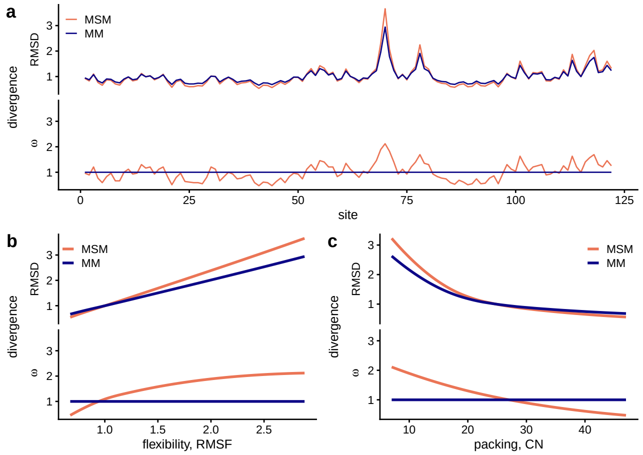


Figure S17: **Comparison of structure and sequence divergence profiles.**

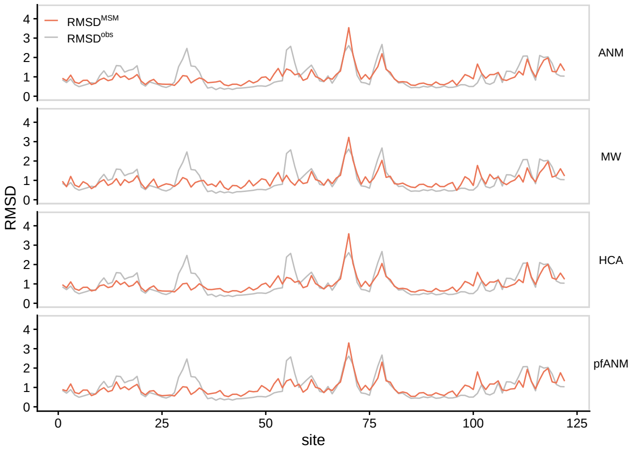


Figure S18: **Dependence of predictions on ENM model choice.**

### RNA recognition motif

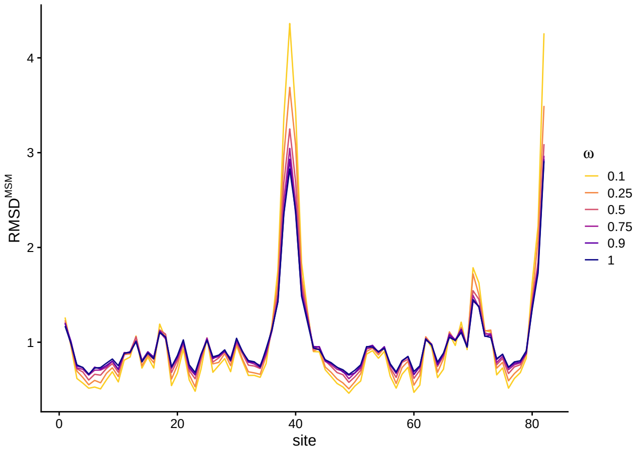


Figure S19: **Predicted site-dependent structure divergence profiles.**

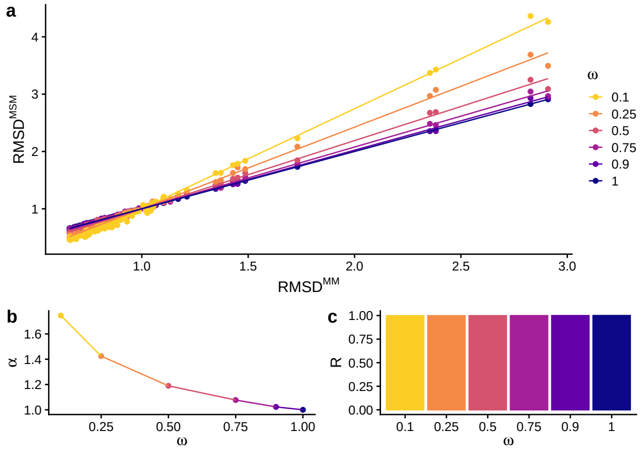


Figure S20: **Structure divergence is proportional to mutational sensitivity.**

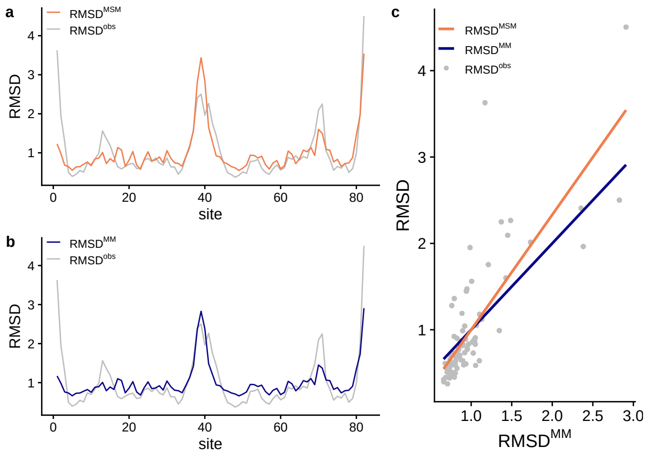


Figure S21: **Predicted vs. observed structure divergence profiles.**

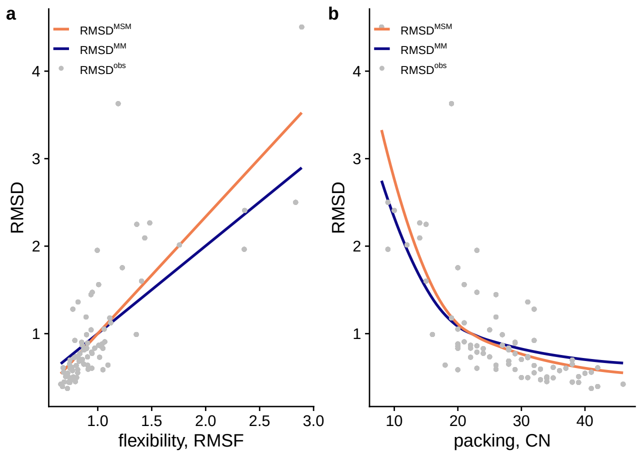


Figure S22: **Dependence of RMSD on flexibility and packing.**

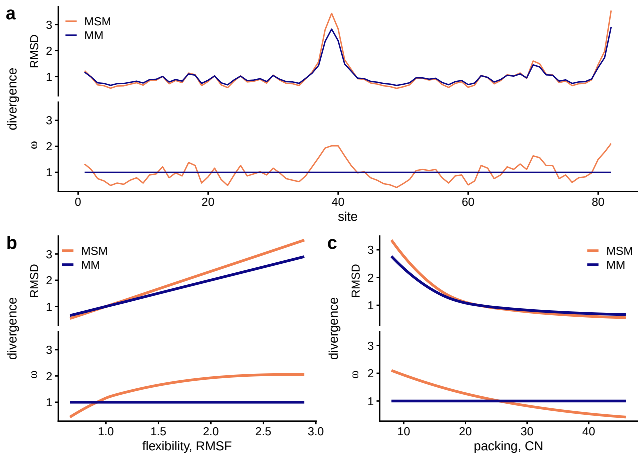


Figure S23: **Comparison of structure and sequence divergence profiles.**

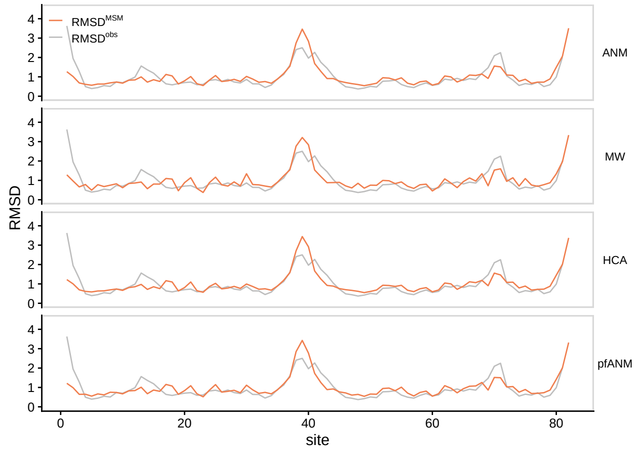


Figure S24: **Dependence of predictions on ENM model choice.**

### Src homology 3 domain

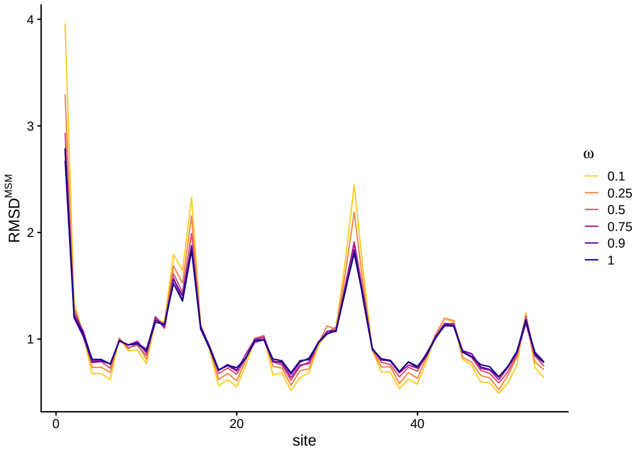


Figure S25: **Predicted site-dependent structure divergence profiles.**

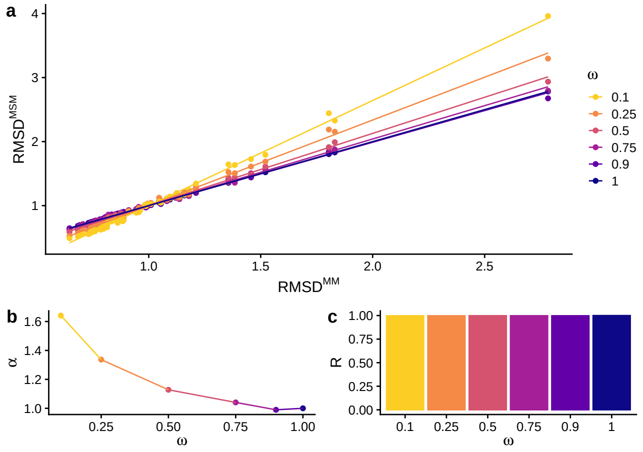


Figure S26: **Structure divergence is proportional to mutational sensitivity.**

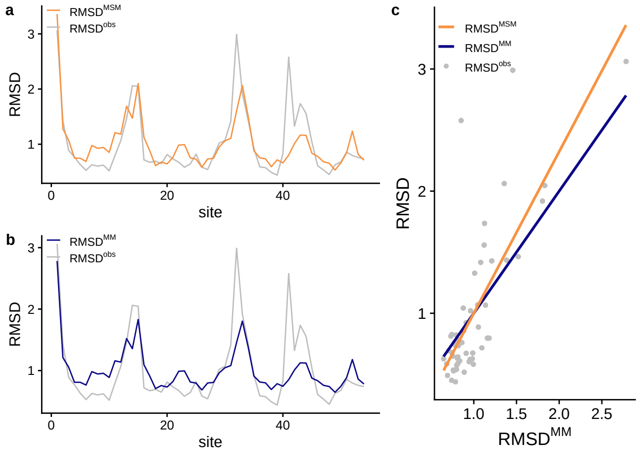


Figure S27: **Predicted vs. observed structure divergence profiles.**

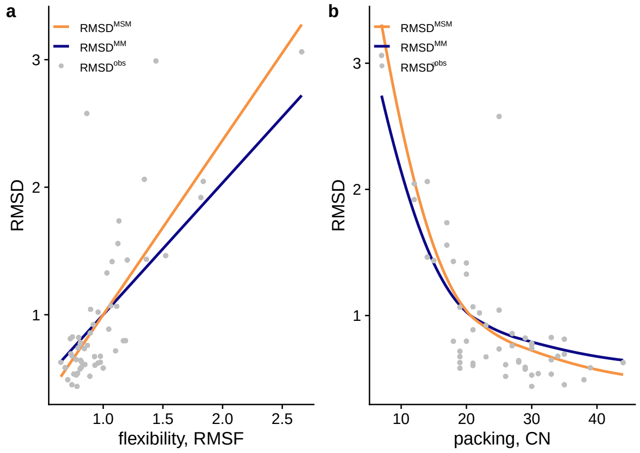


Figure S28: **Dependence of `RMSD` on flexibility and packing.**

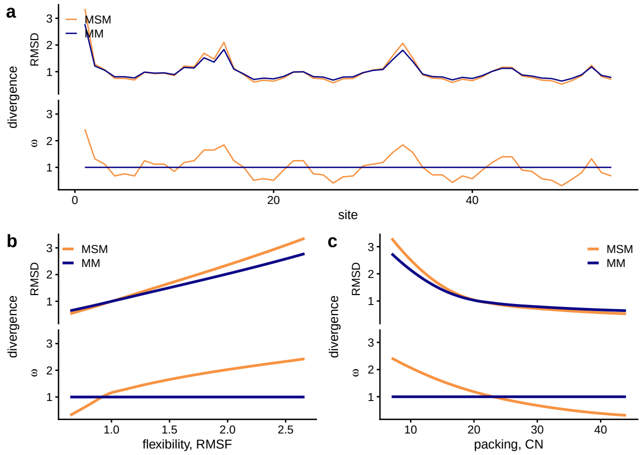


Figure S29: **Comparison of structure and sequence divergence profiles.**

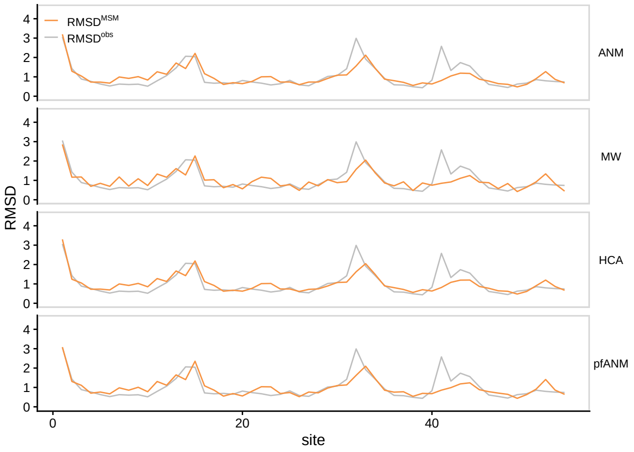


Figure S30: **Dependence of predictions on ENM model choice.**
